# Curability Difference between Autochthonous Mouse Tumors and Their Transplants in Association with Immune Gene Expression

**DOI:** 10.1101/2025.11.23.690042

**Authors:** Hiroshi Tanooka, Chie Kudo-Saito, Fumiko Chiwaki, Masamichi Ishiai, Kouichi Tatsumi, Hiroki Sasaki, Takahiro Ochiya

## Abstract

Compared with transplanted tumors, autochthonous tumors are difficult to cure by experimental radiation therapy in mice. Here we analyzed differences in immune gene expression profiles between mouse fibrosarcomas subcutaneously induced by 3-methylcholanthrene (3MC) and their transplants. The immune genes examined were *pd1, pdl1, pdl2, cd3d, cd8a, cd8b, ifnγ, dx5, grzmb*, and *foxp3*. Among 12 tumors, one was non-transplantable and showed a benign character with an abundance of DX5^+^ natural killer cells and CD8^+^ T cells together with increased IFNγ expression. The other 11 transplantable tumors showed increased expression of *pd1, pdl1, pdl2, cd3d*, and *cd8b*, after their transplantation into syngeneic mice. These effects of transplantation highlight the relevance of immune gene expression status to the curability of tumors.

## INTRODUCTION

Transplanted mouse tumors are widely applied as a model tumor system to experimental radiation therapy [1], while application of autochthonous tumors, such as spontaneous mammary carcinoma [2], is very rare. Autochthonous tumors, compared with transplanted tumors, are difficult to cure by experimental therapy, as noted by Nakahara [3]. According to the strict definition of cure as complete eradication of a tumor mass followed by mouse survival for 120 days without recurrence, fibrosarcomas induced with 3MC in C57BL/6J mice are incurable after experimental radiation therapy with a collimated X-ray beam, with few exceptions, whereas tumors transplanted in syngeneic mice are completely curable in a radiation dose-dependent manner [4] (S1 Fig).

The reason for this distinct difference has long remained unclear. One explanation is that there is new tumor formation at the 3MC-treated site in autochthonous tumors, as it is exposed to the chemical carcinogen 3MC and later to radiation. The recurrent tumors, however, are clones of the original tumors, as demonstrated from analysis of isozyme patterns of tumors in *pgk-1/pgk2* mosaic cell mice [5], indicating true recurrence. One other possible explanation for the curability difference is a change in the immune status of the tumor microenvironment and organismally after transplantation.

Recent findings show that radiation not only directly kills cancer cells, but also induces expression of anti-cancer immune genes [6, 7] and the immunotansmitter [8], and further that the immune activation is more efficient with high LET than low LET radiation [9, 10]. This radiation effect might be related to the transplantation effect for tumor cure.

Here we focused on the expression of immune genes in 3MC-induced mouse tumors before and after transplantation. Target genes were programmed cell death protein (*pd1*) [11], its blockade ligands *pdl1* [12] and *pdl2* [13], the T-cell marker *cd3d* [14], cytotoxic T-cell markers *cd8a* [15] and *cd8b* [15], interferon gamma (*ifnγ*) [16], the perforin *dx5* [17], the integrin *grzmb* [18], and the regulatory T-cell marker *foxp3* [19].

## MATERIALS AND METHODS

### Mice

The experiments were approved by the National Cancer Center (T21-006) and carried out in consideration of the Guideline for Animal Experiments in the National Cancer Center and the 3Rs stipulated by the Animal Welfare Management Act. Germ-free female C57BL/6J mice at 10 weeks of age were purchased from Charles River Japan and kept in the specific pathogen–free room.

### Tumor Induction and Transplantation

An aliquot of 0.1 ml of 3-MC (Sigma) dissolved in 5 mg/ml in olive oil was injected at the groin of mice as described previously [4] . Mice were observed for tumor formation by palpation. When tumors grew to the size of 1 cm in diameter, they were resected and divided into sections for histological examination, transplantation, and gene expression measurements (Fig. 1A). A piece of tumor was transplanted subcutaneously into the groin of a female C57BL/6J mouse with a transplantation needle (13 G) in duplicate.

**Fig 1.**
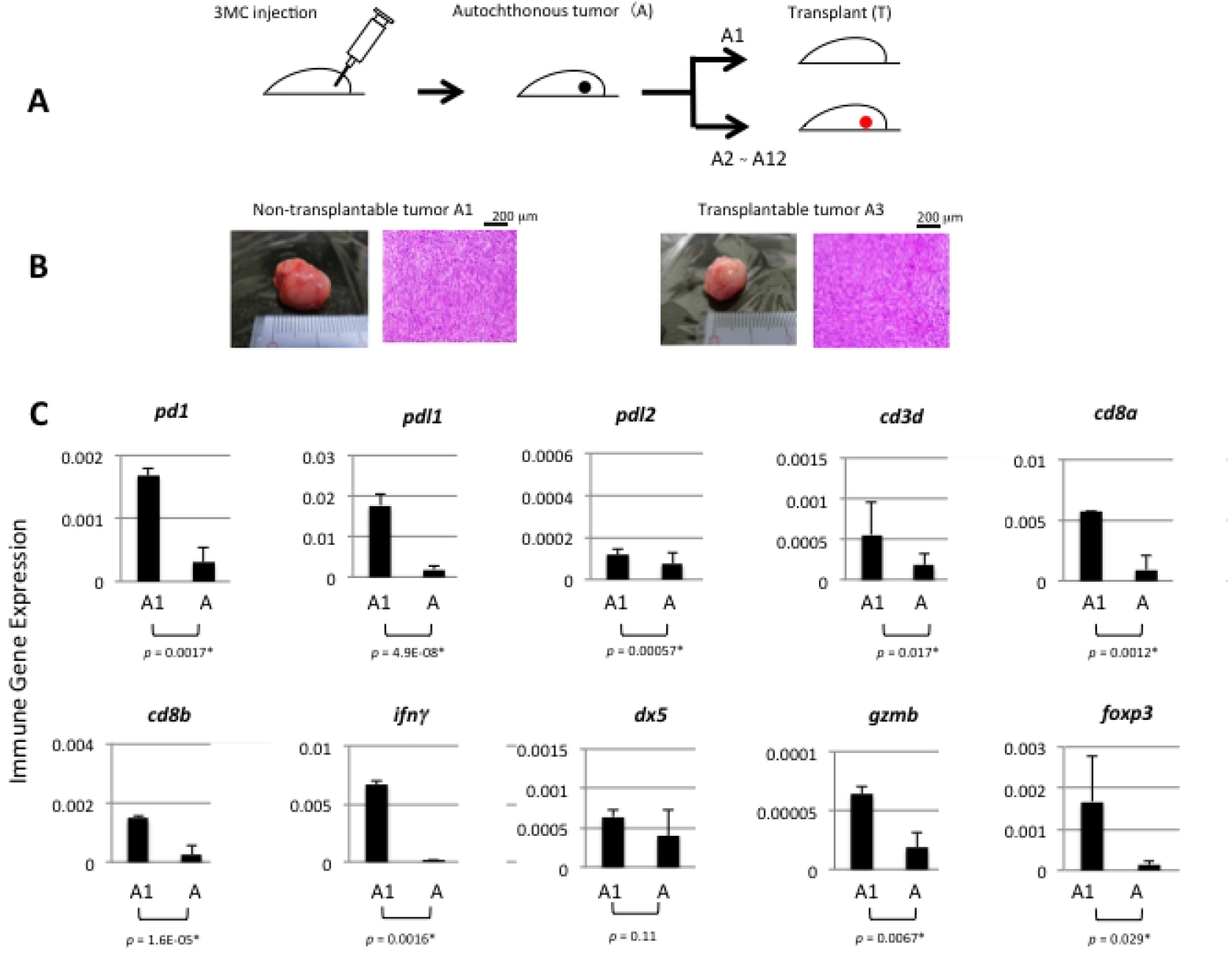
Protocol of experiments, histology and immune gene expression of non-transplantableand transplantable tumors. **A**. Tumor induction with JMC in female C57BL/6j mice and transplantation to syngeneic mice. **B**. Non-transplantable tumor, A1, compared with one of transplantable tumors, A3, showing the indistinguishable histological type offibrosarcomaof the skin. **C**. Expressions ofmRNAs of immune genes in the A1 tumor, measured by qPCR, were compared with the average level of 11 transplantable tumors, A(• : statistically significant, p < 0.05).

Expanded transplants were examined in the same way as for autochthonous tumors. Liver and lung specimens were obtained at the same time as tumor resection. Tumors and organs were kept frozen at -80° C.

### Histology

Tumor sections were fixed in 10% formalin, embedded in paraffin, sliced, stained with hematoxylin and eosin, and examined histologically under a microscope.

### Immune Gene Expression Measurement with qPCR

Two sections of each tumor were homogenized in a lysis reagent (QIAzol, Qiagen) with a homogenizer (Kinematica), and total RNA was extracted with an extraction kit (miRNeasy, Qiagen) and converted to cDNA with a reverse transcription kit (High Capacity cDNA Transcription kit, Applied Biosystems), according to the manufacturer’s protocol. qPCR was performed at 95° C for 3 min followed by 40 cycles at 95° C for 10 s and at 55° C for 30 s with the Real-Time qPCR System (CFX90,Bio-Rad), using a reaction mixture (TaqMan Fast Advanced Master Mix, Applied Biosystems) with probes for measurements of immune gene expression, according to the manufacturer’s protocol (TaqMan Gene Expression Assays): *pd1* (Mm01285676_ml), *pdl1* (Mm03043248_ml), *pdl2* (Mm00451734_ml), *cd3d* (Mm00442746_ml), *cd8a* (Mm01182107_gl), *cd8b* (Mm00438116_ml),*ifnγ* (Mm01168134_ml), *dx5* (Mm00434371_ml), *grzmb* (Mm00493152_gl), *foxp3* (Mm0047516_ml), and *β-actin* (Mm00607939_sl) as a control. Expression levels for immune genes were analyzed using Excel and calibrated to those for *β-actin*.

### Immunohistochemical Analysis

The paraffin-embedded tumor sections were stained with the following antibodies according to standard methods [20]: anti-DX5-FITC (BioLegend #108909), anti-CD8-PE-Cy5 (BD Bioscience #553034), anti-IFNγ-PE (BD Bioscience #554412), and the appropriate isotype controls (BD Bioscience). Three locations per section were observed at 100× magnification using a LSM700 laser scanning confocal microscope (Carl Zeiss), and the number of DX5^+^ NK cells and CD8^+^ T cells contained in the field was counted. The immunofluorescence intensity of IFNγ expression was automatically measured as pixel counts at three locations per section using the ZEN 2012 software installed in the LSM700 microscope. The average of the data was used in graphs.

### Statistics

Student’s t-test was applied to statistical analysis of measured data, using Excel program.

## RESULTS AND DISCUSSION

Autochthonous tumors induced subcutaneously with 3MC in female C57BL/6J mice were compared to transplants for immune gene expression in the context of differences in curability after experimental radiation therapy. Among 13 mice, 12 animals produced tumors (designated as A1–A12). All were fibrosarcomas of the skin and histologically indistinguishable and were transplanted into syngeneic female mice (Fig 1A).

### High infiltration of anti-tumor effector cells in a non-transplantable tumor

Among 12 tumors, one tumor (A1) failed to engraft after four attempts. The other 11 tumors (A2–A12) were transplantable. The A1 tumor was not visibly different from the other transplantable tumors (see Fig 1B for an example comparison). After qPCR analysis of mRNA/cDNA, A1 showed the unique feature of highly expressing *pd1, pdl1, pll2, cd3d, cd8a, cd8b, ifnγ, gzmb*, and *foxp3* as compared to transplantable tumors (Fig 1C). Indeed, immunohistochemical analysis comparing A1 with transplantable tumor tissues highlighted a significantly greater abundance of DX5^+^ natural killer (NK) cells (Fig 2A) and CD3^+^CD8^+^ T cells (Fig 2B) and a significantly higher expression of IFNγ in the infiltrating CD8^+^ T cells (Fig 2C) in the A1 tumor. Of note, a few CD8^+^ T cells but no NK cells were observed within the transplantable tumors, and NK cells in the A1 tumor were much larger than the CD8^+^ T cells (Fig 2D), suggesting an activated state. These results indicate that these anti-tumor effector cells, particularly NK cells, were involved in the benign character of the A1 tumor.

**Fig 2.**
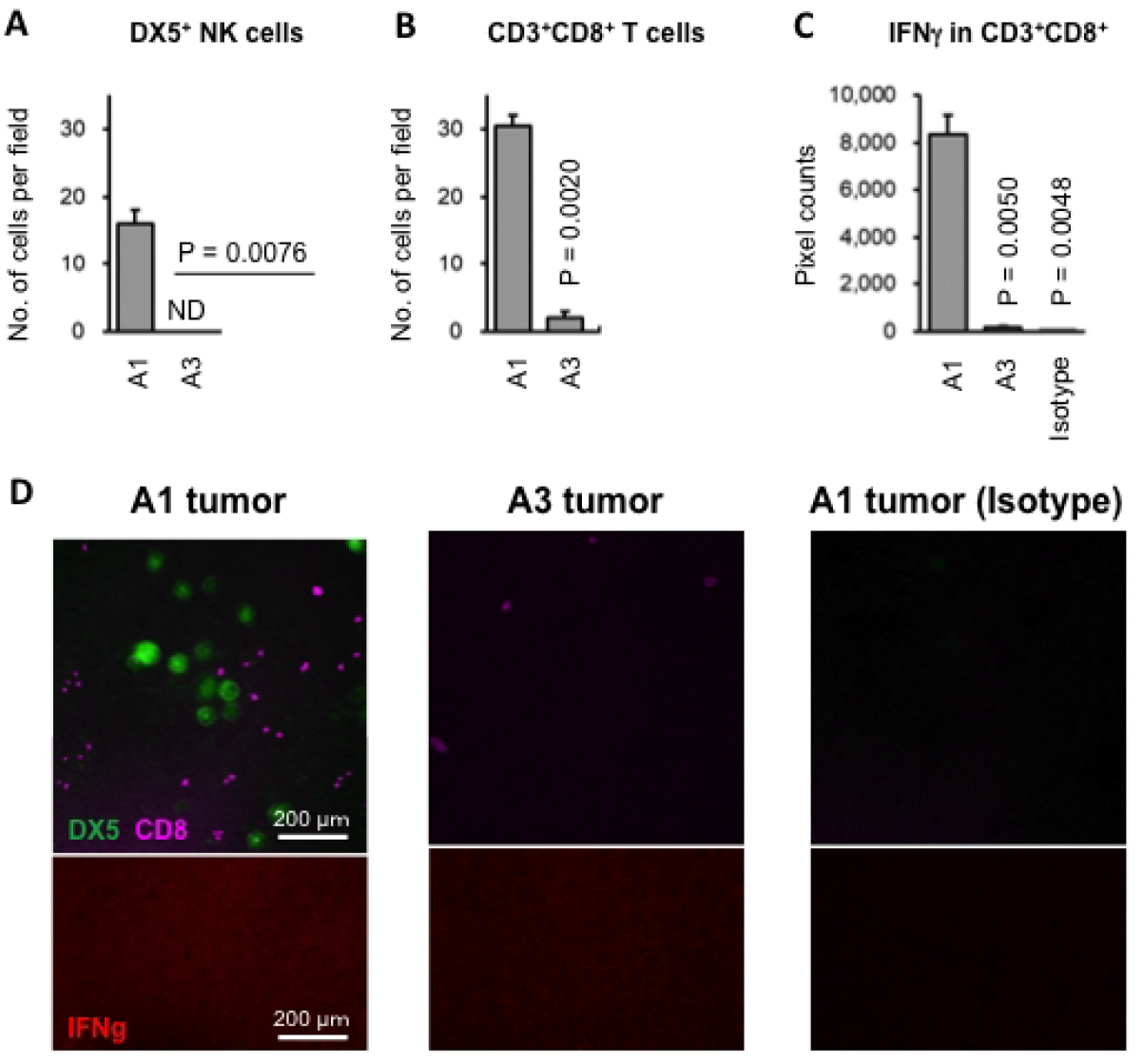
High infiltration of anti-tumor effector cells in non-transplantable tumor. Sections of non-transplantable tumor A1and transplantable tumor A3 were stained with anti-DX5-FITC, anti-CD8-PE-Cy5, anti-lFNγ-PE, and the appropriate isotype controls,. **A**. Infiltration of DX5^+^ NK cells. ND, not detected. **B**. Infiltration of CD8^+^ T cells. **C.** Intensity of lFNγ expression. **D**. Representative photos of A1 and A3 tumors after immuno-histochemical staining.

In a previous study, a few 3MC-induced autochthonous tumors (3 of 28; 11%) were curable after experimental radiation therapy [4]. This frequency largely coincides with non-transplantable tumor frequency in the current study (1 of 12; 8%), indicating that benign tumors can be present in a histologically indistinguishable tumor population even when induced in the same syngeneic mouse strain by the same method.

In the liver of the A1-bearing host mouse, immune gene expression levels did not differ from that in livers of mice bearing transplantable tumors or livers of untreated control mice, while while the lung of A1-bearing mouse showed a higher expression of *pd1, pdl1, pdl2, ifnγ, dx5*, compared with the lung of mice carrying transplantable tumors (S2 Fig). However, this level was at the same level as in untreated control mice, suggesting the possibility that the immune response is suppressed in the 3MC-treated mice carrying transplantable tumors, as shown in the early work [21].

### Immune gene expression in autochthonous tumors and transplants

3MC-induced autochthonous tumors were followed for measurement of immune gene expression after their transplantation into syngeneic mice. The expression levels of *pd1, pdl1, pdl2, cd3d*, and *cd8b* were significantly increased with transplantation (Fig 3). This increase indicates activation of the immune status in tumors and at least partly explains the curability change of autochthonous tumors after transplantation.

**Fig 3.**
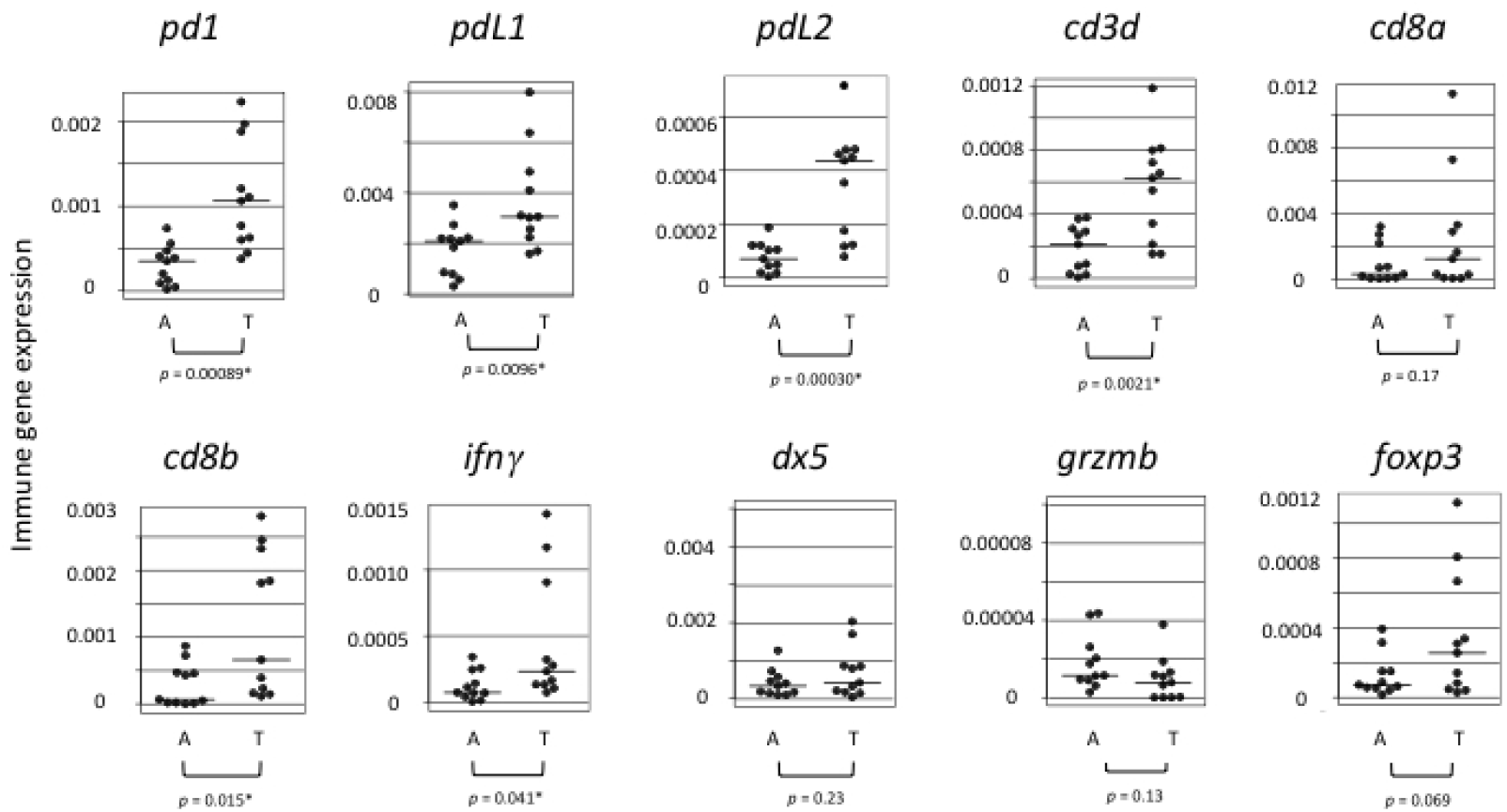
Immune gene mRNA expression in autochthonous tumors and their transplants measured by qPCR. Transplantable 3MC-induced autochthonous fibrosarcomas, A (n = 11), were compared with their transplants, T (n =11), for immune gene expression. Average expression levels are shown by horizontal bars.(•: statistically significant, *p* < 0.05).

Dispersion of the measured values in Fig 3 is thought to reflect the inhomogeneous clonal nature of autochthonous tumors. However, the malignant nature and the immune-enhancing effect are thought to be common in transplantable autochthonous tumors. Cells dying either by transplantation or by irradiation may send similar immune signals into the tumor microenvironment and enhance infiltration of cytotoxic T cells into tumor. The benign tumor, A1 in this study, seems to deliver similar immune signals.

Whether the response of the immune system to radiation is different between autochthonous tumors and their transplants or not is an intriguing problem for the future study,

Additional evidence presents a complex picture. Early studies of the immunogenic nature of 3MC-induced mouse tumors indicated that the host mouse had sufficient immune activity to reject the replanted tumor, whereas untreated mice did not [22, 23], indicating a high anti-tumor immune activity in host animals. Although curability and the immune gene expression profile of tumors were not described in these studies, the findings indicate that host mice bearing autochthonous tumors have a high immunogenic activity and more curable tumors compared with transplanted tumors. This inference, however, is not in keeping with later findings of incurable autochthonous tumors [4].

## CONCLUSION

Immune gene expression in 3MC-induced autochthonous tumors was increased after transplantation. This increase offers possible explanation for the curability difference between autochthonous tumors and their transplants. The benign character of tumor strongly correlated to elevated immune gene expressions.

## Supporting Information

S1 Fig. Curability difference between 3MC-induced autochthonous tumors and their transplants.

S2 Fig. Immune gene expressions in the mouse liver and lung as measured by qPCR.

### Acknowledgments

We thank the late W. Nahakahara and the late R. Tokuzen for advice regarding the 3MC injection method; T. Sado, K. Shimotohno, and H. Nishikawa for useful suggestions and discussions; R. Machinami and A. Yoshida for histological examination; Y. Yamamoto for help in constructing figures; and N. Uchiya and Y. Shiotani for preparing histological sections.

## Data Availability

Data are available upon request to the corresponding author.

